# Reproducible inference of transcription factor footprints in ATAC-seq and DNase-seq datasets via protocol-specific bias modeling

**DOI:** 10.1101/284364

**Authors:** Aslihan Karabacak Calviello, Antje Hirsekorn, Ricardo Wurmus, Dilmurat Yusuf, Uwe Ohler

## Abstract

DNase-seq and ATAC-seq are broadly used methods to assay open chromatin regions genome-wide. The single nucleotide resolution of DNase-seq has been further exploited to infer transcription factor binding sites (TFBS) in regulatory regions via footprinting. Recent studies have demonstrated the sequence bias of DNase I and its adverse effects on footprinting efficiency. However, footprinting and the impact of sequence bias have not been extensively studied for ATAC-seq. Here, we undertake a systematic comparison of the two methods and show that a modification to the ATAC-seq protocol increases its yield and its agreement with DNase-seq data from the same cell line. We demonstrate that the two methods have distinct sequence biases and correct for these protocol-specific biases when performing footprinting. Despite differences in footprint shapes, the locations of the inferred footprints in ATAC-seq and DNase-seq are largely concordant. However, the protocol-specific sequence biases in conjunction with the sequence content of TFBSs impacts the discrimination of footprint from background, which leads to one method outperforming the other for some TFs. Finally, we address the depth required for reproducible identification of open chromatin regions and TF footprints.

## INTRODUCTION

The discovery and characterization of cis-regulatory elements (CREs) such as promoters, enhancers and insulators, is instrumental in delineating the mechanisms of transcriptional gene regulation. These tissue- and developmental stage-specific regulatory elements reside in nucleosome-free, accessible regions of the genome, that are hypersensitive to nuclease attack^1^. Digestion with the nuclease DNase I, coupled to high throughput sequencing (DNase-seq), was the first established genomic technique to probe such open chromatin regions^2,3^, and was widely applied in research consortia such as ENCODE^4,5^ or the Roadmap Epigenomics^6^. A more recent technique, the assay for transposase-accessible chromatin using sequencing (ATAC-seq), employs Tn5 transposase enzymes that preferentially fragment and tag open regions^7^. Both protocols determine genome-wide chromatin accessibility and can locate distal and proximal CREs.

Transcription factors (TFs) bound at CREs are major regulators of gene expression^8^. As protein bound DNA is more resistant to cleavage with DNase I, leaving behind protected stretches of nucleotides or shortly “footprints”^9^, DNase-seq potentiates the inference of TF-bound locations genome-wide (TF-footprinting)^10,11^. A multitude of TF-footprinting methods have been developed to date^12^, which can be grouped under three general categories: site-centric, segmentation based, and integrative site-centric methods. Site-centric methods model footprints specifically for candidate TF binding sites (TFBSs), using the shape or magnitude of the DNase-seq signal around them^13–16^. Segmentation based methods, on the other hand, scan the DNase-seq signal for footprint-like signatures (eg. peak-trough-peak pattern) and subsequently match the identified footprints to putative TFs^17–21^. Integrative site-centric methods model bound sites using combinations of diverse features, such as motif match score, sequence conservation and variable length bins of DNase-seq signal around candidate TFBSs^22–27^.

The efforts to assay bound sites genome-wide via TF-footprinting have come under scrutiny by studies demonstrating that DNase I cleaves the underlying DNA in a non-uniform manner, where sequence composition dictates the cleavage propensities (also known as sequence bias)^28,29^. This necessitates the discrimination of actual footprints from footprint-like signal profiles originating solely due to sequence bias^16^. To account for this, a number of TF-footprinting tools explicitly model and incorporate the bias background in their models or processing pipelines, by calculating the ratio of observed to expected DNase cuts for short sequences of fixed length^12,15,20^. 6-mers have been the primary choice, as they capture enough variation to represent the bias^16^, in line with the finding that the main sequence information content around a DNase cut site is confined to the flanking 3 nucleotides on either side^28^. Open chromatin regions^12,16,20^ or DNase-seq experiments conducted on deproteinized genomic DNA^12,15^ have been used to infer these 6-mer cleavage propensities.

Recent efforts have explored the feasibility of TF-footprinting with ATAC-seq^24,25,30^, however this is not yet studied as extensively as for DNase-seq. Furthermore, like DNase I, Tn5 transposase is reported to have specific sequence preferences^30,31^, but the effect of this on ATAC-seq TF-footprinting efficiency is not systematically investigated. It is thus unclear whether the same set of sites would be identified as footprints using ATAC-seq and DNase-seq in a comparative setting. Here, we infer footprints using data obtained from DNase-seq and a modified ATAC-seq protocol in the same cell line, taking the enzyme-specific sequence biases into account, and we show that despite the difference in footprint shapes, the locations identified as bound are in concordance. We report that TF-footprinting efficiency is closely linked to clear discrimination of the footprint from the background, which is dependent on the enzyme-specific biases and the sequence content of the TFBSs, making one method more preferable than the other for some TFs. We also address the largely open question on library depth that is required for identification of open chromatin regions and footprints, based on the irreproducible discovery rate (IDR) in conjunction with libraries sequenced to different depths. Our analysis demonstrates that careful consideration of the inherent sequence bias and assessment of reproducibility render TF-footprinting feasible, even at moderate sequencing depths.

## METHODS

### DNase-seq and ATAC-seq experimental procedures and data preprocessing

DNase-seq and ATAC-seq assays were performed on human cell lines, K562 and HEK293 cells. K562 and HEK293 cells were cultured in Iscove’s Modified Dulbecco’s Medium (IMDM) and Dulbecco’s Modified Eagle’s Medium (DMEM), respectively, both complemented with 10% fetal bovine serum (FBS) and 1% Penicillin/Streptomycin.

DNase-seq experiments were conducted on 50 million cells as previously described^32^, with the minor modification of using 5’ phosphorylated oligo 1b. Samples digested with 12U, 4U and 1.2U total DNase I were pooled. Libraries constructed from pooled digests were sequenced on the Illumina HiSeq2500 platform using the single-end sequencing mode with 50-bp reads. Analysis was conducted in line with the official ENCODE DNase-seq pipeline. Specifically, the reads were trimmed to the first 20 bases, as only this portion corresponded to the ends of DNase I-digested fragments, due to the MmeI cleavage step in the protocol. Trimmed reads were aligned to the hg19 build of the human genome, using the Burrows-Wheeler aligner (BWA)^33^, tolerating up to two mismatches. Sequences aligning to more than four locations were discarded. Further processing was performed to filter out unwanted chromosomes and problematic regions such as alpha satellites. In order to remove PCR artifacts, reads that piled up (>=10 reads) at a single base were discarded, if they constituted at least 70 percent of all reads in the surrounding 30 base pair window.

ATAC-seq experiments were performed on 50000 cells for the K562 samples and 100000 cells for the HEK293 samples, following the published protocol^7^ but increasing transposition time from 30 minutes to 1 hour for all samples. In addition, lysis conditions were varied in different experiments. For the K562 sample denoted “10 minute lysis”, cell lysis was performed via a 10 minute centrifugation in lysis buffer, as described in the original protocol^7^. For the K562 sample denoted “5 minute lysis”, a shorter lysis of 5 minutes was used. For the K562 sample denoted “no lysis buffer” and all HEK293 samples, the centrifugation in lysis buffer step was omitted altogether, and the cell pellets were taken directly to the transposition reaction. Libraries were sequenced on the Illumina HiSeq2000 platform, with 100-bp paired end reads. Since fragments as short as 38 base pairs were expected, adapter sequences were trimmed from the 3’ end of the reads. Specifically, matches of any length to the reverse-complemented Nextera Transposase Adapters (CTGTCTCTTATACACATCTGACGCTGCCGACGA, CTGTCTCTTATACACATCTCCGAGCCCACGAGAC) were removed. Trimmed reads were aligned to the hg19 build of the human genome, using bowtie2^34^ with parameter -X set to 1500, to allow correct alignment of paired-end fragments up to 1500 base pairs. Only reads that aligned uniquely to a single location were retained, by filtering out the multimappers marked with the XS:i flag in the sam file. PCR duplicates were removed using Picard (http://broadinstitute.github.io/picard/). Further processing was performed to filter out contigs as well as the Y and mitochondrial chromosomes, and retain only reads that aligned concordantly as a pair within the expected fragment length range (38-1500 bp).

Library complexity and saturation were calculated using the preseq program^35^, using the c_curve and lc_extrap functionalities. Correlations of reads counts between libraries were calculated using the bamCorrelate bins command of the deepTools suite, with the parameters –corMethod pearson, -bs 100, --fragmentLength 1 and –doNotExtendPairedEnds.

### Peak calling

In order to find open chromatin regions, peak calling was performed on the processed DNase-seq and ATAC-seq datasets using JAMM^36^, with parameters -f 1 and -d y. Parameter -f 1 ensured taking only the 5’ ends of the reads into account which corresponded to the actual cleavage/transposition sites. As duplicates were already removed prior to peak calling, parameter -d y was used to keep all processed reads.

Where replicates were available, peaks in agreement between the two replicates were found using the irreproducible discovery rate (IDR) pipeline^37^. Specifically, the “batch-consistency-analysis.r” script of the pipeline was executed using the “signal.value” parameter, ranking the peaks of the two replicates by signal intensity for comparison. The “half.width” and “overlap.ratio” parameters were set to -1 and 0, respectively, where true peak widths were used without alteration and two peaks were considered to be part of the same region if there was at least 1bp overlap between them. The number of peaks that were found to be concordant at the stringent 0.01 IDR threshold was noted. Then, JAMM was once again used, this time to call peaks on the two replicates together rather than individually, with the -f 1,1 parameter. In this way, peaks were called where both replicates consistently displayed signal enrichment. This peak set was further truncated using the number obtained from the IDR analysis, resulting in the final JAMM-IDR peaks.

### Sequence bias of Tn5 transposase

The sequence bias of the Tn5 transposase was calculated in the form of 6-mers, similar to the previous calculations of DNase bias^15^. To this end, libraries generated by Tn5 transposition on deproteinized genomic DNA (see supplementary table 2) were preprocessed in the same way as ATAC-seq datasets as detailed above. As the 5’ ends of the reads corresponded to the transposition sites, the sequences of all 6-mers centered on these sites were retrieved (eg transposition between the third and fourth nucleotides). Occurrences of all these 6-mers in the data were counted and the relative frequencies were calculated for each. Similarly, background genomic frequencies were calculated by counting all 6-mers present in the mappable portion of the genome. The frequencies observed in the data were normalized to the background frequencies to obtain the final transposition propensities per 6-mer. Deviations from one indicated increased or decreased propensities, thus bias.

The average Tn5 transposition propensity in a candidate binding site of a given transcription factor was calculated by retrieving and counting all 6-mers associated with the site (without flanks). The counts were multiplied by the Tn5 transposition propensities of the associated 6-mers, summed and normalized by the total number of 6-mers in the site. The same calculation was applied for DNase, using the previously calculated DNase cleavage propensities per 6-mer^15^.

### Scanning the genome for candidate binding sites

The SpeakerScan Toolset^38^ was used to scan the hg19 build of the human genome with position weight matrices (PWMs), to find candidate transcription factor binding sites (TFBS). PWMs contain expected frequencies for each nucleotide in a per-base fashion, modeling the binding sequence preferences of a given TF. A pseudocount of 0.0005 was added to each frequency in the PWMs, to ensure non-zero entries. At each PWM-sized window in the genome, a TFBS score was calculated, as the log-likelihood of the underlying sequence matching the PWM versus a background model. The background was modeled with a first order Markov chain in a 500 bp local window, centered on the considered position. The top scoring 50000 sites were taken along for transcription factor footprinting in this study.

### Identification of transcription factor footprints

Transcription factor footprinting was performed with a site-centric method from our lab as previously described^15^. Specifically, candidate TFBSs were considered with 25bp flanks upstream and downstream (parameter PadLen=25). Parameter k=2 was used to model two components; one for the footprint and one for the background. Both components were modeled as multinomials along the considered window size (TFBS+50bps), where each value corresponded to the cleavage/transposition probabilities at a given nucleotide. For the footprint component, these probabilities were found by computing the aggregate DNase or ATAC-seq signal (from the 5’ ends of the reads) around the TFBSs that overlap ChIP-seq peaks for that factor and re-estimating the signal via expectation maximization. For the background component, the probabilities were calculated as the signal that would be expected solely due to the protocol-specific bias values, given the sequences around the candidate TFBSs (parameter Background=“Seq”). As we had previously not observed a distinct difference in performance, the background was kept fixed and not re-estimated (parameter Fixed=T). Once both components were learned, footprint scores were calculated for all candidate TFBSs, as the log-odds of footprint versus background (footprint log-likelihood ratio, FLR).

The IDR strategy was applied here as well where replicates were available, to find reproducible footprints. To this end, candidate TFBSs with positive FLRs in both replicates were chosen and ranked by FLR. IDR analysis was performed with the same parameters as explained for peak calling, where FLR values replaced signal intensities. Again, the number of sites that passed the stringent 0.01 IDR threshold was noted. Finally, TFBSs were ranked by the average FLR from the two replicates and truncated according to the IDR result. This led to the reproducible FLR-IDR footprints.

Footprint model AUCs were calculated by 4-fold cross validation. Briefly, the data was split into 4 parts, and for TFBSs in each part, FLR was calculated using footprint and background models learned from the other 3 parts. TFBSs were ranked by FLR, and those intersecting ChIP-seq peaks were labeled as the true positives. The AUCs obtained from the four parts were averaged to obtain the final value.

Our footprinting pipeline for the Linux command line can be downloaded from https://ohlerlab.mdc-berlin.de/software/Reproducible_footprinting_139/. A Galaxy implementation of our code is also available at https://toolshed.g2.bx.psu.edu/view/rnateam/footprint, with the associated Conda package at https://bioconda.github.io/recipes/footprint/README.html.

## RESULTS

### A modified ATAC-seq protocol decreases mtDNA contamination and improves agreement with DNase-seq

Early ATAC-seq libraries generated with the original protocol have large numbers of reads mapping to mitochondrial DNA (mtDNA) that need to be discarded, which severely impacts the final library depth^7^. For an ATAC-seq library where we followed this protocol, we made the same observation in K562 cells, with 75% of the reads mapping to mtDNA (figure 1A supplementary table 1). To decrease the mtDNA contamination, we evaluated two different approaches: decreasing the time of cell lysis to 5 minutes in lysis buffer (from the original 10 minutes) and eliminating the lysis buffer step altogether by proceeding directly to the transposition reaction. Of these, particularly the approach where no lysis buffer was used, led to a substantial improvement, with only 18% percent of the reads mapping to mtDNA in this library (figure 1A supplementary table 1), in line with previous reports^39^. Avoiding the detergent lysis may help mitochondrial membranes to stay intact, with other forces such as osmotic pressure being adequate to permeabilize the nuclear membrane.

**Figure 1:**
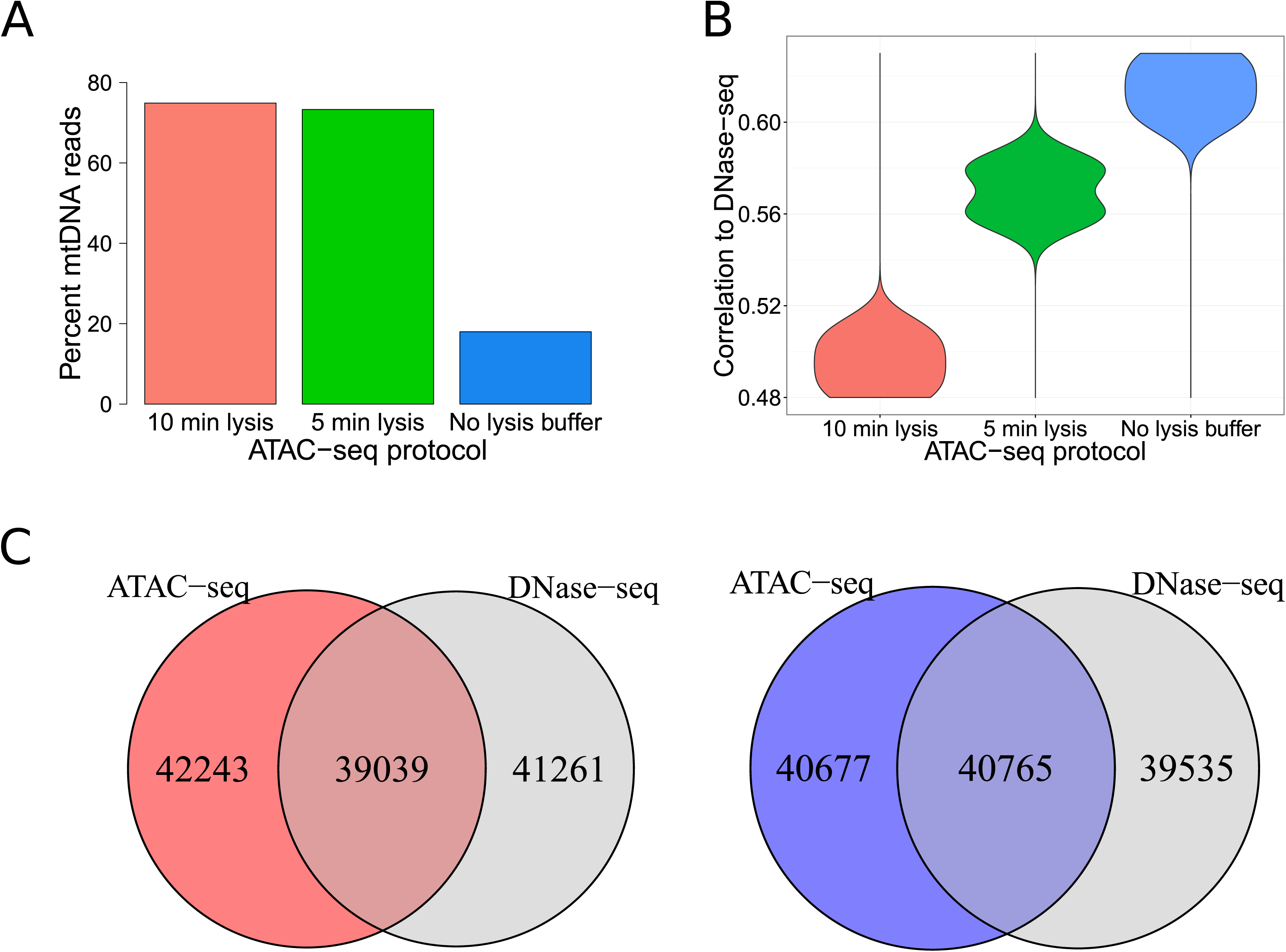
Generating ATAC-seq libraries without the usage of lysis buffer increases agreement with DNase-seq. (A) Percentage of all reads that align to the mitochondrial genome in K562 ATAC-seq libraries generated with the published protocol (10 min lysis), shorter lysis (5 min lysis) or without using lysis buffer (no lysis buffer). (B) Agreement of these libraries with all K562 DNase-seq libraries as measured by Pearson correlations of read counts in 100bp bins genome wide. (C) Overlap of peaks found in K562 DNase-seq data with peaks in ATAC-seq data generated using the published protocol (left) and peaks in ATAC-seq data generated without using lysis buffer (right).

To adequately quantify the protocol-related differences of ATAC-seq vs. DNase-seq, we also generated a single-hit DNase-seq library in K562 cells, and compared this alongside three other publicly available single-hit DNase-seq datasets (supplementary table 2) with the ATAC-seq libraries. Avoiding the usage of lysis buffer also increased the read-level agreement between the two experimental approaches (figure 1B Pearson correlations of read counts in 100 base pair bins; supplementary figure 1). This effect was already partially visible in data from the short lysis protocol. To investigate whether this observation is also reflected at the region-level of open chromatin, we called peaks with JAMM^36^ and identified the set of concordant peaks using the irreproducible discovery rate (IDR) pipeline for DNase-seq data where replicates were available (see methods)^37^. Using the peak signal values for ranking, at the stringent 0.01 IDR threshold, we found 80,300 JAMM-IDR peaks for DNase-seq. We selected the same number of top scoring peaks in the ATAC-seq datasets, since replicates were not available for these libraries. The overlap between DNase-seq peaks and ATAC-seq peaks obtained with the original protocol was 39,039 (48.6%) (figure 1C left) and increased to 40,765 (50.8%) for the ATAC-seq protocol where lysis buffer was not used (figure 1C right). This improved agreement at the open chromatin region-level, albeit moderate, provided further support that avoiding detergent lysis increases the concordance between ATAC-seq and DNase-seq.

### Open chromatin regions are found reliably at moderate library depths

The library depth of next-generation sequencing protocols that is required for a given downstream application is not always clear, especially when the regions of interest are not as clearly defined as e.g. protein-coding genes. To investigate the effect of library depth on uncovering open chromatin regions, we generated 11 ATAC-seq libraries with different depths in HEK293 cells using the protocol with no lysis buffer (four high, three medium and four low-depth libraries, figure 2A and supplementary table 1). The individual libraries were derived from two biological replicates. To obtain the highest possible depth representing these two samples (>300,000,000 read pairs each), all technical replicates were merged and denoted by “combined ATAC-seq replicates”. Alongside the ATAC-seq experiments, we generated a single-hit DNase-seq library in HEK293 cells and additionally downloaded and processed two publicly available single-hit DNase-seq replicates (supplementary table 2). We observed strong positive correlations between all ATAC-seq and DNase-seq libraries at the level of genome-wide read counts (0.62-0.77 Pearson correlations of read counts in 100 base pair bins; supplementary figure 2), and JAMM-IDR peaks called for the combined ATAC-seq and DNase-seq replicates showed again a significant overlap (supplementary figure 3A).

**Figure 2:**
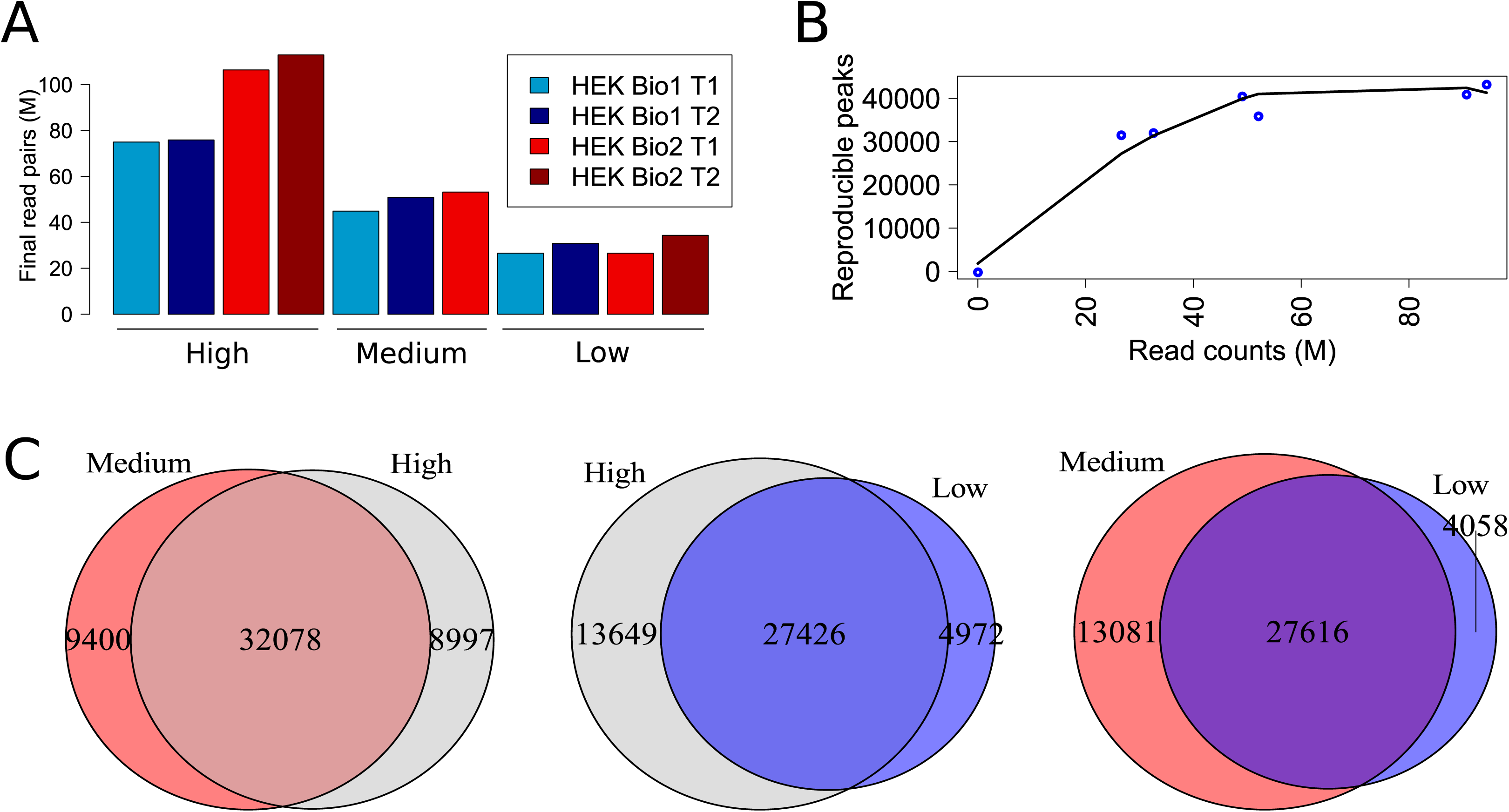
The task of finding open chromatin regions saturates at medium depth. (A) Number of reads after processing in the 11 HEK293 ATAC-seq libraries with different library depths. The two biological replicates are shown in blue and red, with the shades representing the technical replicates. (B) Numbers of reproducible peaks found with the JAMM-IDR strategy at different depths. (C) The overlaps between one set of peaks in (B) shown for high vs. medium (left), high vs. low (middle) and medium vs. low sets (right).

**Figure 3:**
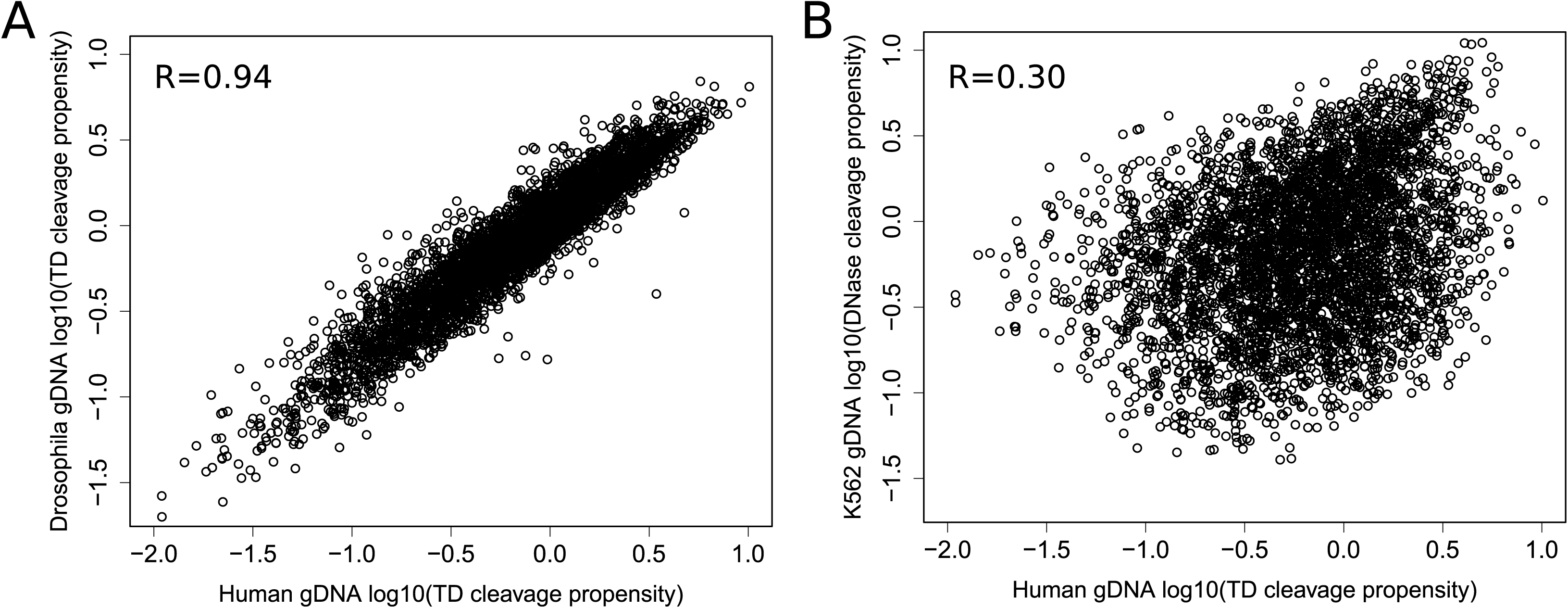
The sequence bias of the Tn5 transposase is distinct from that of DNase I. (A) Comparison of Tn5 transposition propensities of all 6-mers (log10 scale) in two libraries generated using deproteinized genomic DNA from human (YH1) and D.melanogaster. (B) 6-mer transposition propensities in the human library compared to cleavage propensities of DNase inferred previously from a single-hit DNase-seq experiment using deproteinized genomic DNA from K562 cells.

We then investigated to what extent the individual ATAC-seq libraries sequenced at different depths could capture the open chromatin regions uncovered by the combined replicates. To this end, libraries of similar depth from different biological replicates were matched in a pairwise manner to get JAMM-IDR peaks (supplementary table 3). This resulted in six total peak sets, corresponding to two of each of high, medium and low-depth library comparisons. Similar numbers of peaks were found at high and medium depth, with a slight decrease at low depth (figure 2B supplementary figure 3A). Additionally, these peak sets displayed notable agreement among themselves and with the peaks of the combined ATAC-seq dataset (figure 2C supplementary figure 3A). These observations suggested near-saturation for the task of defining open chromatin regions, even though none of the libraries were at saturation at these depths (supplementary figure 4). Moreover, these six IDR peak sets showed 63% to 72% overlap with the peaks of the DNase-seq data, which exceeded the 61% observed for the combined ATAC-seq data (supplementary figure 3A); even though a higher number of peaks was found in the combined dataset, IDR analysis of the individual datasets led to more reproducible subsets of the total pool. In support of this, the peaks found in the combined ATAC-seq dataset that did not overlap any of the peaks in the six individual sets, were predominantly low-signal, distal regions (supplementary figure 3B). Taken together, replicate libraries of low to medium depth of 25-50 million reads were sufficient for reliable identification of open chromatin regions in human cell lines.

**Figure 4:**
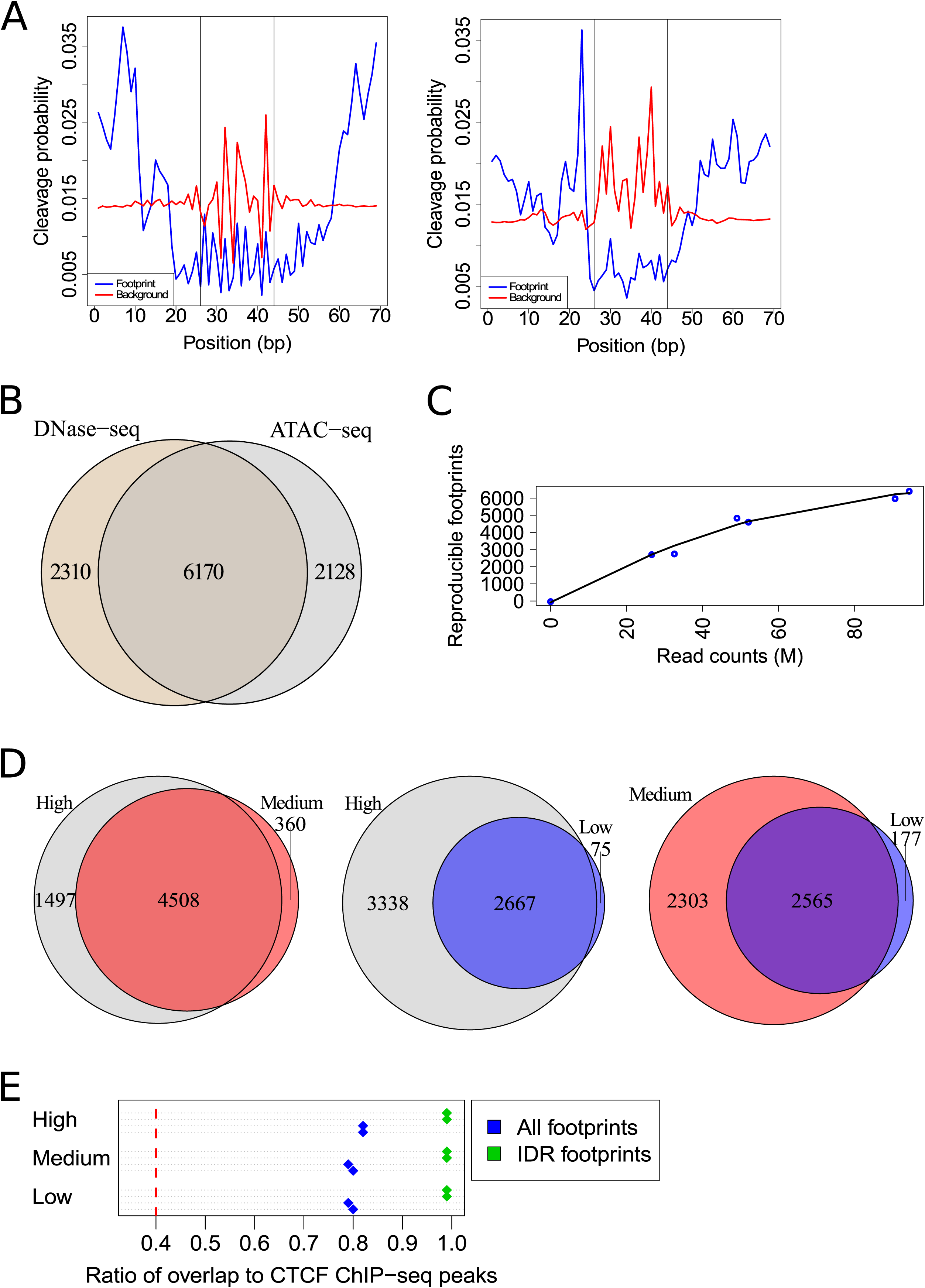
The number of reproducible footprints scales with library depth. (A) CTCF footprints inferred from HEK293 ATAC-seq data (left) and DNase-seq data (right). Vertical lines depict the edges of the motif match. (B) Overlap between reproducible CTCF footprints in the HEK293 DNase-seq and combined ATAC-seq replicates, found using the FLR-IDR strategy. (C) Numbers of reproducible CTCF footprints in HEK293 ATAC-seq datasets at different depths. (D) The overlaps between one set of footprints in (C) shown for high vs. medium (left), high vs. low (middle) and medium vs. low sets (right). (E) The ratio of reproducible CTCF footprints (IDR footprints) or all CTCF motif regions with positive footprint scores (all footprints) that overlap CTCF ChIP-seq peaks, in all six individual sets at different depths (supplementary table 3). Red dashed line indicates this ratio for all considered CTCF motif sites.

### Sequence bias of ATAC-seq deviates from that of DNase-seq

A multitude of studies have explored the efficacy of transcription factor footprinting with DNase-seq. These studies have demonstrated that the DNase I enzyme cleaves genomic DNA in a non-random fashion, where it has different cut propensities for different sequences, and this sequence bias has adverse effects on the quality of footprinting when left uncorrected^16^. Our lab has previously published a site-centric computational footprinting tool where 6-mer DNase bias has been incorporated into the model to estimate the bias background in a multinomial mixture framework^15^. In order to gain insights into the sequence bias of ATAC-seq data, we calculated the 6-mer cleavage propensities of the Tn5 transposase, using available data from libraries generated by Tn5 transposition on deproteinized genomic DNA^31^ (supplementary table 2). Comparison of the cleavage propensities in libraries generated using human genomic DNA vs. *D*.*melanogaster* genomic DNA, revealed very similar results (figure 3A Pearson correlation 0.94), indicating that the Tn5 transposase has specific sequence preferences which are consistent in data from the two species. The dynamic range of this bias is on the same order of magnitude as for DNase bias^15^. We next asked how the sequence preferences of the Tn5 transposase compare to those of DNase I. Using values inferred previously from a single-hit DNase-seq experiment of deproteinized K562 cells^15^, we observed this correlation to be fairly low (figure 3B Pearson correlation 0.30). This indicated that these enzymes have largely distinct sequence biases.

### ATAC-seq and DNase-seq generate different footprint shapes for the same factor

In order to systematically examine how ATAC-seq compares to the more established DNase-seq method in transcription factor footprinting, we first focused on CCCTC binding factor (CTCF), a factor with a well-known, high information content binding site with substantial available ChIP-seq data including in HEK293 cells (supplementary table 4). We scanned the human genome for matches to the CTCF binding model obtained from the JASPAR database (supplementary table 5). As aggregate signal across all candidate CTCF motif matches is expected to be a mixture of footprint (bound sites) and background (unbound sites), our method^15^ was applied to infer the bound subset by modeling the shapes of the CTCF footprints in the DNase-seq and combined ATAC-seq replicates. Shape of the aggregate signal at sites that overlap CTCF ChIP-seq peaks was used to initialize the footprint model. The background was modeled using protocol-specific bias values. The resulting footprint and background profiles revealed marked differences between ATAC-seq and DNase-seq (figure 4A left and right, respectively). Most notable was a wider region of protection in the ATAC-seq data, in line with a previous study^31^ which reported that the Tn5 transposase dimer needs circa 30 nucleotides to bind DNA and that cleavage occurs in the central 9 nucleotides. Another difference concerned the background profiles, attributable to the distinct sequence preferences of these two enzymes. In short, from the same set of CTCF motif matches, different footprint and background models were learned using ATAC-seq and DNase-seq datasets.

### Footprinting using ATAC-seq and DNase-seq uncovers common bound sites

This observation led to the question whether the same sites would be identified as bound by a transcription factor when using ATAC-seq and DNase-seq in the same cell type. Using the protocol-specific footprint and background models learned for CTCF, we calculated the footprint scores for all considered motif matches, as the log-odds of footprint versus background per site (footprint log-likelihood ratio, FLR, see methods). The FLR is thus derived in a protocol-specific manner, solely from the single-nucleotide resolution signal around motif sites, without relying on additional features, and it accounts for sequence bias, making it an ideal metric to compare the footprints from the two protocols. As a positive FLR indicates a higher probability of being bound vs. unbound, we selected the motif matches that had a positive FLR in both replicates of the assayed method. We again used IDR to find the reproducible subset of CTCF footprints among these sites, ranked by FLR (FLR-IDR, see methods). Following this methodology for the combined ATAC-seq replicates, 12,651 motif sites had positive FLRs in both replicates, of which 8,298 were found to be reproducible by FLR-IDR (supplementary figure 5). For the DNase-seq replicates, of the 13,592 sites with positive FLRs, 8,480 were reproducible. Nearly all of the reproducible footprints of ATAC-seq and DNase-seq overlapped CTCF ChIP-seq peaks (98% and 96% respectively, supplementary figure 5). Furthermore, these reproducible footprints from the two experimental protocols were also concordant, with 6,170 sites (74%) overlapping (figure 4B). This analysis of ATAC-seq and DNase-seq data thus identified many common sites as bound, despite the difference in footprint shapes.

**Figure 5:**
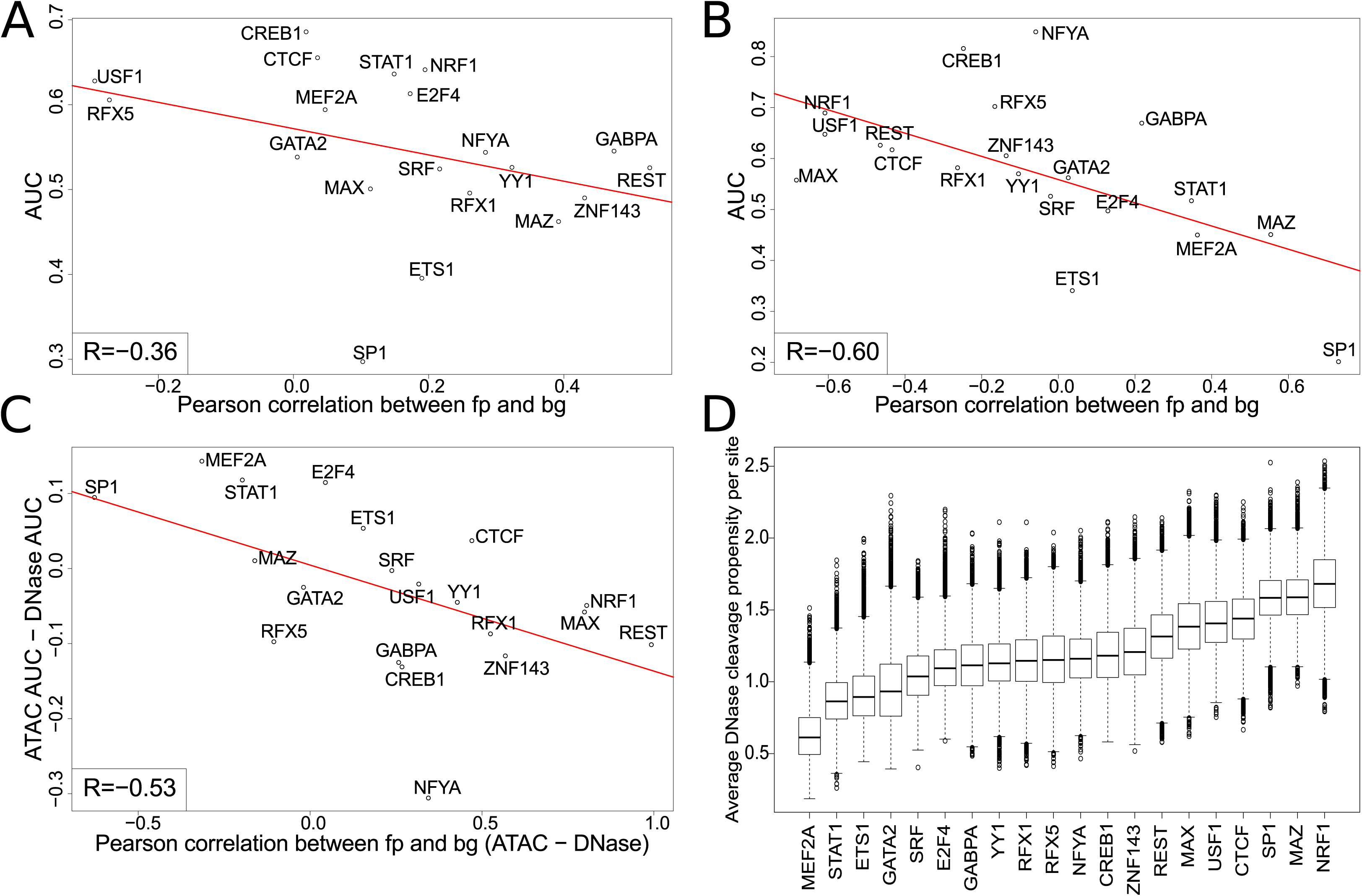
TF-footprinting accuracy is linked to clear discrimination of footprint from background. (A,B) AUCs vs footprint-background model similarities in (A) ATAC-seq data and (B) DNase-seq data. (C) Difference in AUCs (ATAC-DNase) vs difference in footprint-background model similarities (ATAC-DNase). (D) Average DNase I cleavage propensities over candidate TFBSs for all 20 assayed factors.

### Number of reproducible footprints scales with library depth

Previous studies that inferred cell-type specific TF binding site annotations from DNase footprint data typically used very large datasets (with hundreds of millions of reads per cell type)^10,17^. To investigate the feasibility of footprinting at lower library depths, we next conducted the analysis on the 11 individual ATAC-seq libraries. We used the same setup for pairwise comparisons as for peak calling (supplementary table 3), this time to find reproducible CTCF footprints at different library depths. Even though the numbers of motif matches that had positive footprint scores were in the same range for all analyzed pairs, the numbers of reproducible footprints gradually declined with decreasing depth (figure 4C supplementary figure 5). This indicated that, unlike peak calling, footprinting efficiency did not saturate and rather followed the library complexities at these depths (supplementary figure 4). However, the footprints at distinct depths had substantial overlaps with each other and also constituted almost perfect subsets of the footprints found in the combined ATAC-seq data (figure 4D supplementary figure 5). Moreover, these reproducible footprint sets consistently showed 99% overlap with CTCF ChIP-seq peaks, compared to around 80% when considering all motif sites with positive FLRs (figure 4E). Taken together, even though deeper sequencing is beneficial to footprinting coverage, the assessment of reproducibility enables finding smaller but equally reliable sets of footprints at lower depths.

### The properties of the footprints generalize to multiple transcription factors

To elucidate whether the previous observations would also apply more generally beyond CTCF, we conducted the footprinting analysis on other factors. The limited availability of ChIP-seq data in HEK293 cells motivated an experimental setup to learn the footprint shapes in K562 cells, where ChIP-seq data is more abundant (supplementary table 4), and use these models to find footprints in HEK293 cells. To this end, all ATAC-seq data in K562 cells was merged to get adequate depth (supplementary table 1) and among the K562 DNase-seq datasets, the second ENCODE replicate was chosen (supplementary table 2). As proof of principle, we first confirmed that the CTCF footprint shapes were almost identical to those learned from HEK293 data (supplementary figure 6A). We then learned footprint models from K562 data for 19 additional transcription factors with available ChIP-seq data (supplementary tables 4 and 5). For a subset of these factors, namely NRF1, CREB1 and USF1, the footprint shapes reflected the expected protection pattern in both ATAC-seq and DNase-seq data; in line with the previous observations from CTCF motif regions, the ATAC-seq footprints displayed a wider region of protection compared to the DNase-seq footprints (shown for NRF1 in supplementary figure 7A). The footprint scores (FLR) for these three factors and CTCF were in close correspondence with the associated ChIP-seq signal values in K562 cells, conferring further confidence in these footprint models (supplementary figure 6B-E). Thus, we used these models to identify bound sites reproducibly with the FLR-IDR strategy in HEK293 cells. As for CTCF, reproducible footprints were found to be concordant between DNase-seq and combined ATAC-seq replicates; at the level of individual HEK293 ATAC-seq datasets, library depth and the numbers of reproducible footprints showed again a strong dependency (shown for NRF1 in supplementary figure 7B and C, respectively). As the observations could be replicated for multiple factors, these results likely provide insights into the general properties of the footprints.

### Protocol-specific sequence biases influence footprinting efficiency

Strong footprints that were concordant in both ATAC-seq and DNase-seq data were only found for four of the 20 assayed factors. For most factors, clear footprints were observed in one of the experimental methods, but not the other. Therefore we asked whether the distinct sequence biases of the two methods play a role in the factor-dependent performance of footprinting. To get a continuous measure for performance (as opposed to the discrete visual assessment of footprint shapes), for all TFs in both experimental settings, we calculated the area under the receiver operating characteristic curve (AUC), ranking candidate sites by FLR and considering those that overlap ChIP-seq peaks to be true binding sites. In order to assess how performance is linked to the relationship between the footprint and background models, the Pearson correlations between these two models (eg. footprint-background model similarities) for each TF were calculated and compared to the AUCs. The AUCs negatively correlated with the footprint-background model similarities in both ATAC-seq and DNase-seq datasets (figure 5A and B, correlations of -0.36 and -0.6, respectively), indicating that when a footprint model is clearly distinguished from the background, it is more likely to explain transcription factor binding accurately. Moreover, the differences per TF between ATAC-seq and DNase-seq datasets for these two measures (AUCs and footprint-background model similarities), also had a negative correlation (−0.53, figure 5C), suggesting that the experimental protocol which achieves better separation between the footprint and background components, is also performing better for a given TF. As the background component is derived directly from sequence bias, we next explored the role of bias more explicitly. In particular, two factors for which ATAC-seq outperformed DNase-seq, MEF2A and STAT1, had the lowest DNase I cleavage propensities (eg. sequence bias) over their motif regions, among all assayed factors (figure 5D), whereas the Tn5 transposition propensities for these factors were average (supplementary figure 8A). Therefore, the background models learned from DNase bias for these factors had footprint-like shapes, impeding the clear separation between the two components, and thus explaining the poor performance of DNase-seq (shown for MEF2A in supplementary figure 8B). In summary, due to the distinct sequence biases of ATAC-seq and DNase-seq, the sequence content of transcription factor binding sites can influence footprinting efficiency in a protocol-specific manner.

## DISCUSSION

DNase-seq has been widely used to assay open chromatin regions and TF footprints. The emergence and increasing use of ATAC-seq, necessitates a systematic comparison of the two methods, especially for TF-footprinting. Here, in a comparative setting, we have shown that although the two methods have distinct sequence biases and generate different footprint shapes for the same TF, the sites they identify as bound are largely in agreement. However, the sequence content of TFBSs combined with protocol-specific sequence biases, impact footprinting efficiency for some TFs, leading to larger differences for these factors and making one method preferable to the other.

There are opposing views on the library depth required for TF-footprinting. Whereas some studies require at least 200 million reads^10^, others demonstrate efficient TF-footprinting at moderate sequencing depths (50-60 million reads)^23,30^, in agreement with our results. These moderate numbers were reported for both segmentation-based^30^ and integrative site-centric^23^ tools, challenging the view that these approaches have different depth requirements^10^. To get the highest possible depth, pooling all replicates has been a common practice in TF-footprinting studies. However, our results indicate that keeping the replicates separate to assess reproducibility may lead to more accurate footprint predictions. This is especially relevant for low-depth libraries, where this approach enables finding reliable subsets of the total footprint pool.

Although the sequence bias of DNase I is well characterized, there is still no consensus about the benefits of bias correction for TF-footprinting. Whereas some studies report increased accuracy upon bias correction^12^, others do not make this observation^23^. One explanation for this might be the different approaches to DNase signal processing and TF-footprinting. Methods that extensively smooth the signal, or use features that diverge from single-nucleotide resolution (eg. binned signal) might be less affected by bias. Since our method has single-nucleotide resolution, we have used protocol-specific biases to model the background in our TF-footprinting approach. As TF-footprinting efficiency depends on the clear separation of footprint from background, ATAC-seq outperformed DNase-seq for two factors with low average DNase I cleavage propensities over their motif regions that resulted in a footprint-like background profile. Surprisingly, the opposite was not as clear to observe, e.g. for factors where DNase-seq outperformed ATAC-seq, the average Tn5 cleavage propensities over the motif regions were not consistently at the lower end of the spectrum. Furthermore, the range of average cleavage propensities over all TFs was narrower for Tn5 (figure 5D vs supplementary figure 8A). This may indicate that 6-mers perform slightly better in modeling the sequence bias of DNase I compared to Tn5 transposase.

Our comparative analysis therefore clearly confirms previous reports that DNase cleavage bias might render footprints of some factors “invisible”, and that, in general, performance to identify footprints can vary significantly across assays and TFs. At the same time, these observations point to a strategy to address this problem: for effective footprinting for all TFs, one could combine assays with different sequence biases. Finally, in contrast to previous studies that reported no correlation between ChIP-seq signal values and footprint scores^20^, we have previously observed and now observe again a strong link between these two measures, implying that the footprint score we have defined here is a quantitative measure of occupancy. In summary, we expect that the insights gained from this work will provide experimental design and computational analysis guidelines for future TF-footprinting studies.

## ACKNOWLEDGEMENTS

This work was partially supported by grants from the German Federal Ministry of Education and Research (BMBF) as part of the RNA Bioinformatics Center of the German Network for Bioinformatics Infrastructure (de.NBI) [031 A538C RBC] as well as the Human Frontier Science Program (HFSP RGY0093/2012). We thank Galip Gürkan Yardimci for feedback on the analysis methods and the manuscript. We thank Rebecca Hunt for feedback on the manuscript and the code.

## AUTHOR CONTRIBUTIONS

A.K.C. and U.O. developed the analysis approach. A.K.C. analyzed the sequencing data with supervision from U.O. A.H. performed the ATAC-seq and DNase-seq experiments. R.W. and A.K.C. implemented the command-line version of the code. D.Y. carried out the Galaxy implementation. A.K.C. and U.O. wrote the manuscript.

