## Supplementary Materials for "Reproducible inference of transcription factor footprints in ATAC-seq and DNase-seq datasets via protocol-specific bias modeling"

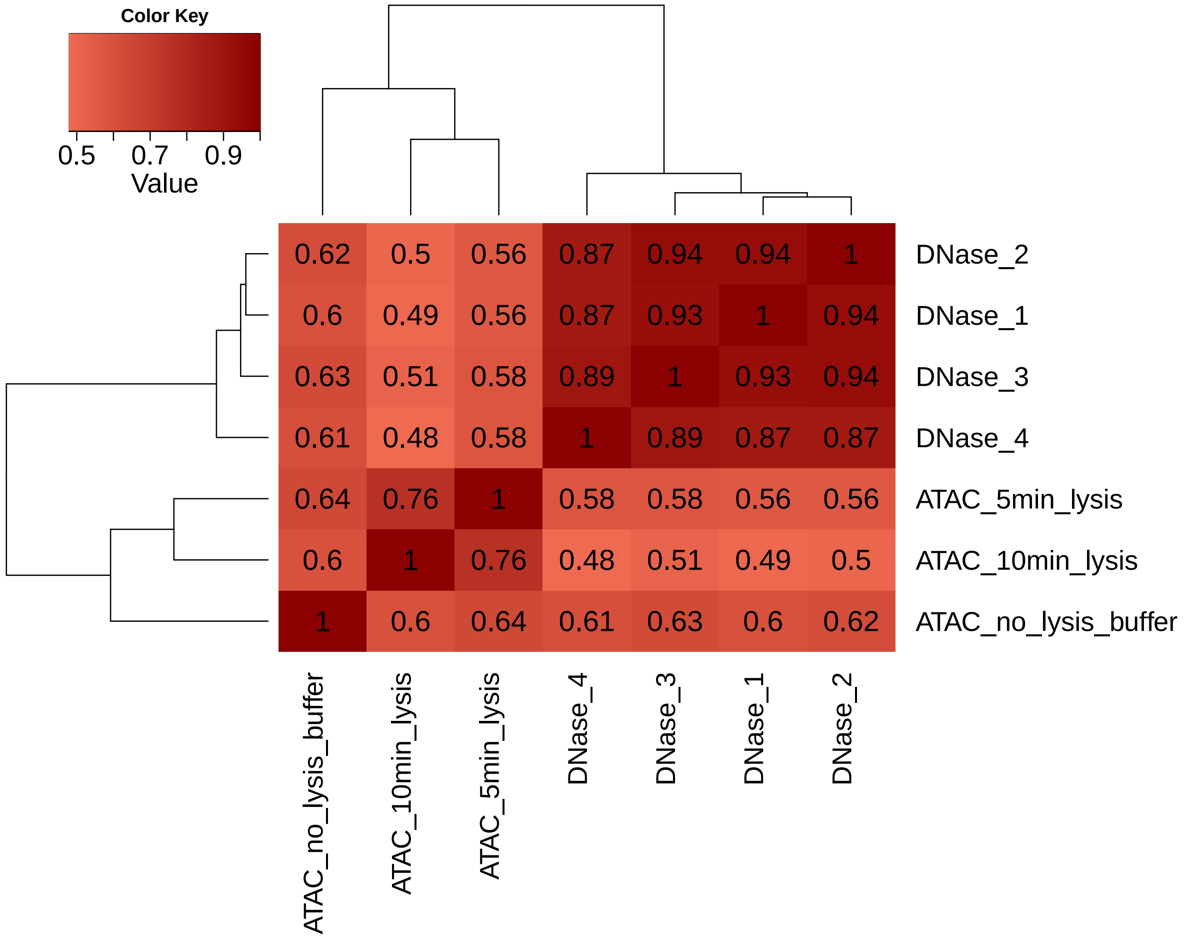


**Supplementary Figure 1:** Pairwise Pearson correlations of read counts in 100bp bins genome-wide for all ATAC-seq and DNase-seq datasets in K562 cells. ATAC-seq datasets are labeled with the employed protocol: 10 min lysis (published protocol), 5 min lysis and no lysis buffer. DNase 1-3 are the replicates from the ENCODE project and 4 is the library newly generated for the study, all following the single-hit protocol.

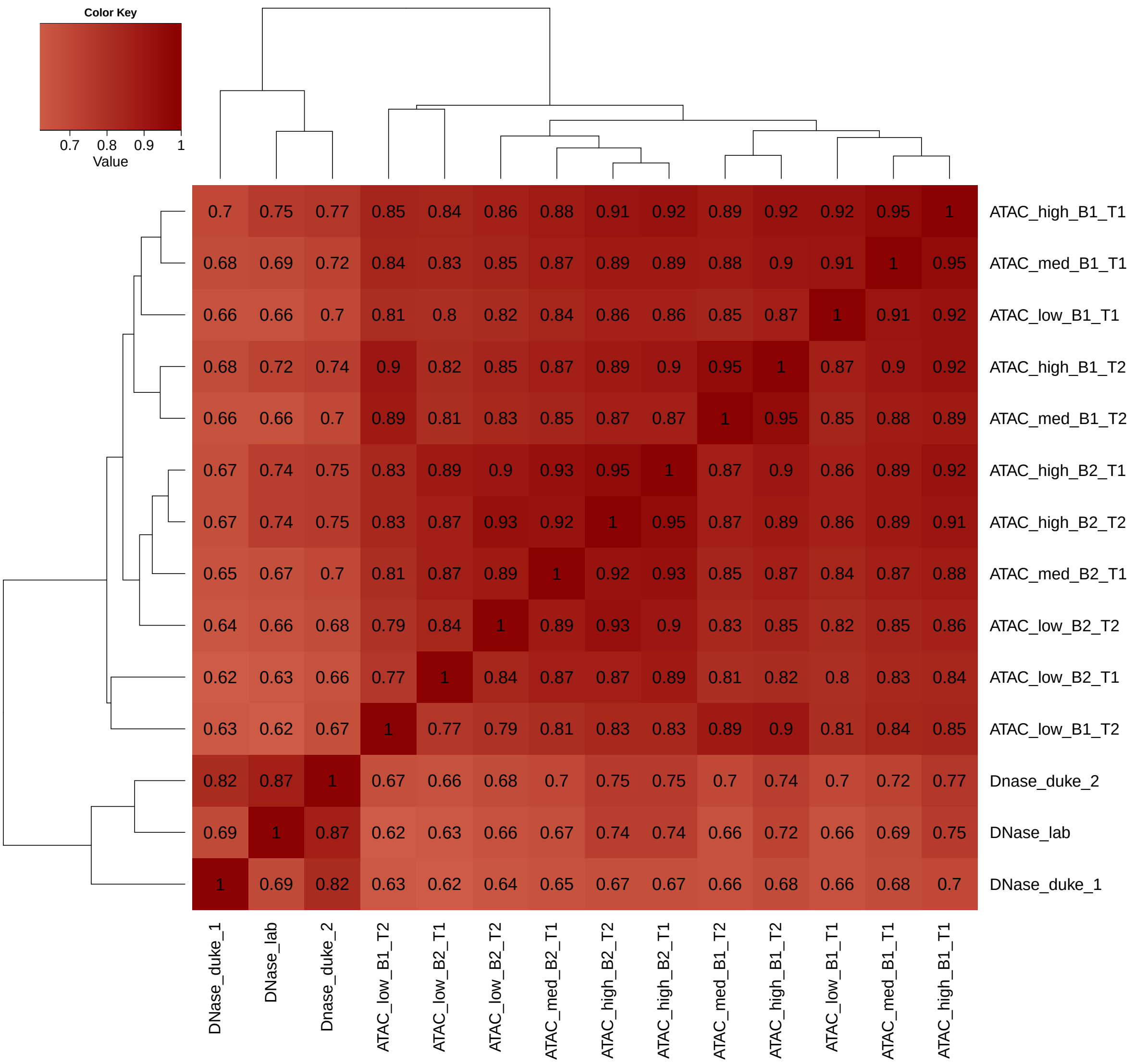


**Supplementary Figure 2:** Pairwise Pearson correlations of read counts in 100bp bins genome-wide for the ATAC-seq and DNase-seq datasets in HEK293 cells. All ATAC-seq datasets are generated with the protocol where no lysis buffer is used. The corresponding library depth (high, medium or low), biological (B1 or B2) and technical (T1 or T2) replicate status is indicated. DNase 1 and 2 are the replicates from the ENCODE project and lab refers to the library newly generated for the study, all following the single-hit protocol.

A

| | DNase | Combined_ATAC | High_ATAC1 | High_ATAC2 | Medium_ATAC1 | Medium_ATAC2 | Low_ATAC1 | Low_ATAC2 | Number of peaks |
|---|---|---|---|---|---|---|---|---|---|
| DNase | | 0.61 | 0.51 | 0.51 | 0.52 | 0.48 | 0.44 | 0.44 | 56019 |
| Combined_ATAC | 0.48 | | 0.52 | 0.53 | 0.53 | 0.47 | 0.43 | 0.43 | 68970 |
| High_ATAC1 | 0.66 | 0.86 | | 0.77 | 0.78 | 0.72 | 0.67 | 0.65 | 41075 |
| High_ATAC2 | 0.63 | 0.84 | 0.74 | | 0.73 | 0.68 | 0.63 | 0.62 | 43381 |
| Medium_ATAC1 | 0.66 | 0.86 | 0.77 | 0.76 | | 0.74 | 0.68 | 0.66 | 40647 |
| Medium_ATAC2 | 0.68 | 0.87 | 0.8 | 0.79 | 0.84 | | 0.73 | 0.72 | 36053 |
| Low_ATAC1 | 0.72 | 0.9 | 0.84 | 0.83 | 0.87 | 0.83 | | 0.77 | 31674 |
| Low_ATAC2 | 0.71 | 0.89 | 0.82 | 0.82 | 0.85 | 0.81 | 0.76 | | 32183 |

B

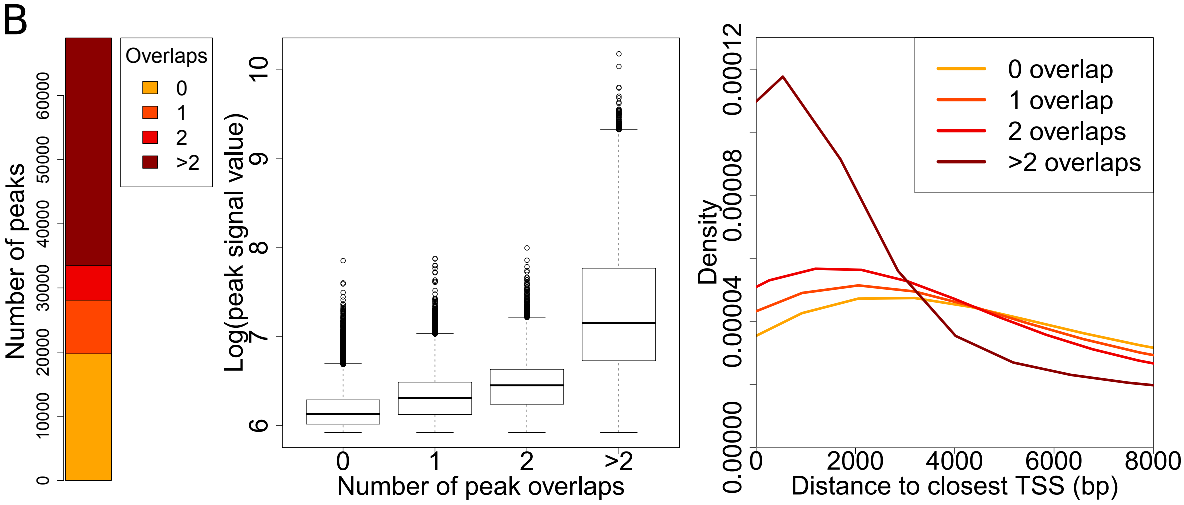


**Supplementary Figure 3:** Analysis of reproducible peaks in HEK293 cells. (A) Overlaps between all reproducible JAMM-IDR peaks found in HEK293 DNase-seq and ATAC-seq datasets. The number in each cell represents the ratio of the peaks in the row-dataset that overlap the peaks of the column-dataset. Total numbers of peaks are given on the right. (B) Number of JAMM-IDR peaks in the combined ATAC-seq replicates that overlap the union of peaks from the six individual datasets zero, one, two or more times (left). Peak signal values (middle) and distance to closest TSS (right) are shown for these four groups.

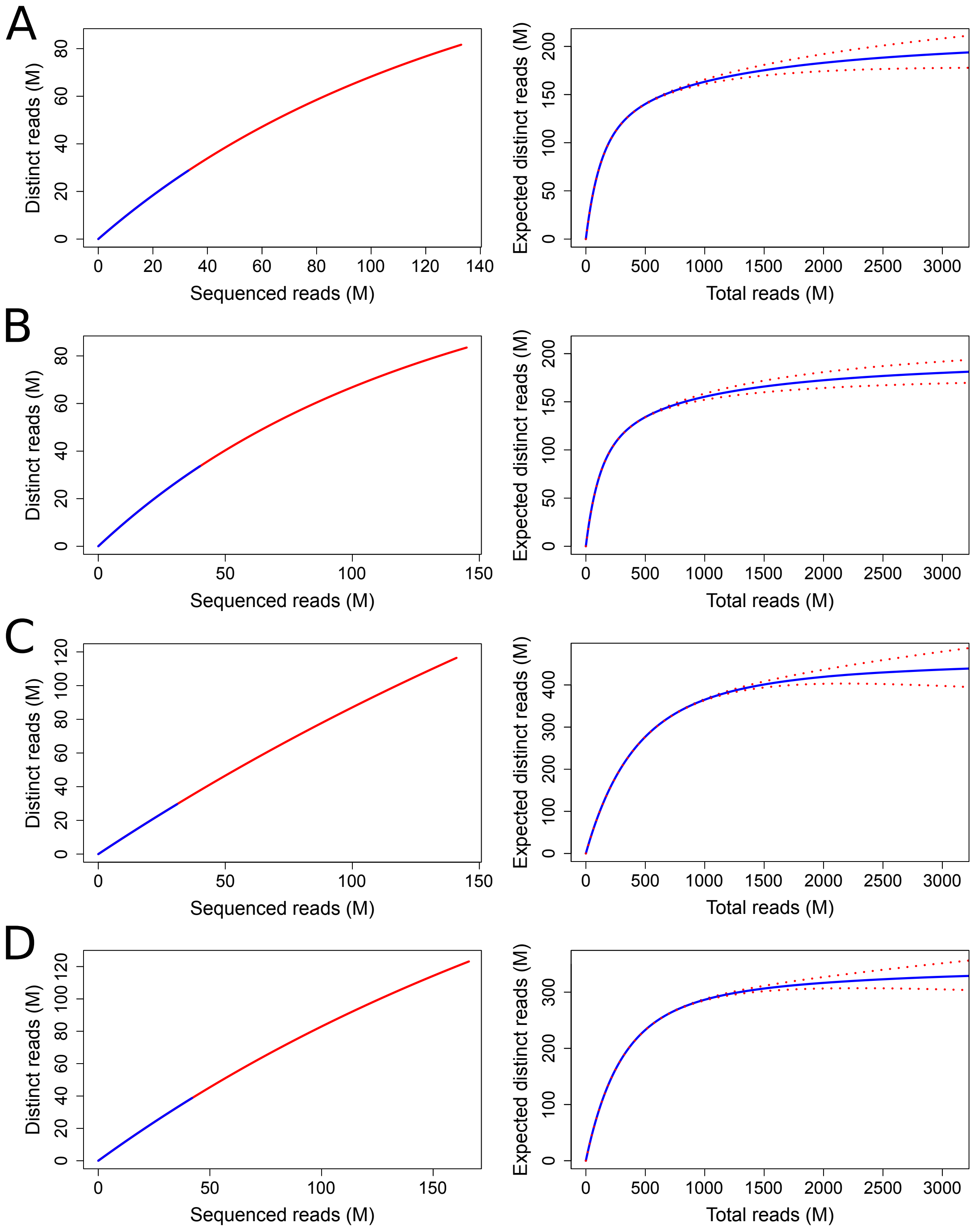


**Supplementary Figure 4:** Library complexity and saturation plots for HEK293 ATAC-seq datasets. (A-D) Complexity (left) and saturation plots (right) for (A) biological replicate 1 technical replicate 1 (B1-T1), (B) B1-T2, (C) B2-T1 and (D) B2-T2. Library complexity is shown at high and low library depth levels, in red and blue, respectively.

| | DNase | Combined_ATAC | High_ATAC1 | High_ATAC2 | Medium_ATAC1 | Medium_ATAC2 | Low_ATAC1 | Low_ATAC2 | Number of reproducible/ total footprints | Overlap with ChIP peaks |
|---|---|---|---|---|---|---|---|---|---|---|
| DNase | | 0.73 | 0.58 | 0.6 | 0.48 | 0.45 | 0.28 | 0.28 | 8480/13592 | 8151 (96%) |
| Combined_ATAC | 0.74 | | 0.72 | 0.76 | 0.58 | 0.55 | 0.33 | 0.33 | 8298/12651 | 8114 (98%) |
| High_ATAC1 | 0.81 | 0.99 | | 0.87 | 0.75 | 0.7 | 0.44 | 0.45 | 6005/12473 | 5938 (99%) |
| High_ATAC2 | 0.8 | 0.99 | 0.82 | | 0.69 | 0.68 | 0.41 | 0.42 | 6435/12308 | 6339 (99%) |
| Medium_ATAC1 | 0.83 | 0.99 | 0.93 | 0.91 | | 0.82 | 0.53 | 0.52 | 4868/12204 | 4818 (99%) |
| Medium_ATAC2 | 0.83 | 0.99 | 0.91 | 0.94 | 0.87 | | 0.53 | 0.55 | 4634/12107 | 4585 (99%) |
| Low_ATAC1 | 0.86 | 1 | 0.97 | 0.96 | 0.94 | 0.9 | | 0.68 | 2742/11225 | 2718 (99%) |
| Low_ATAC2 | 0.87 | 1 | 0.96 | 0.98 | 0.91 | 0.91 | 0.67 | | 2782/11591 | 2768 (99%) |

**Supplementary Figure 5:** Overlaps between all reproducible FLR-IDR CTCF footprints found in HEK293 DNase-seq and ATAC-seq datasets. The number in each cell represents the ratio of the footprints in the row-dataset that overlap the footprints of the column-dataset. Numbers of footprints and their overlaps with ChIP-seq peaks are given on the right.

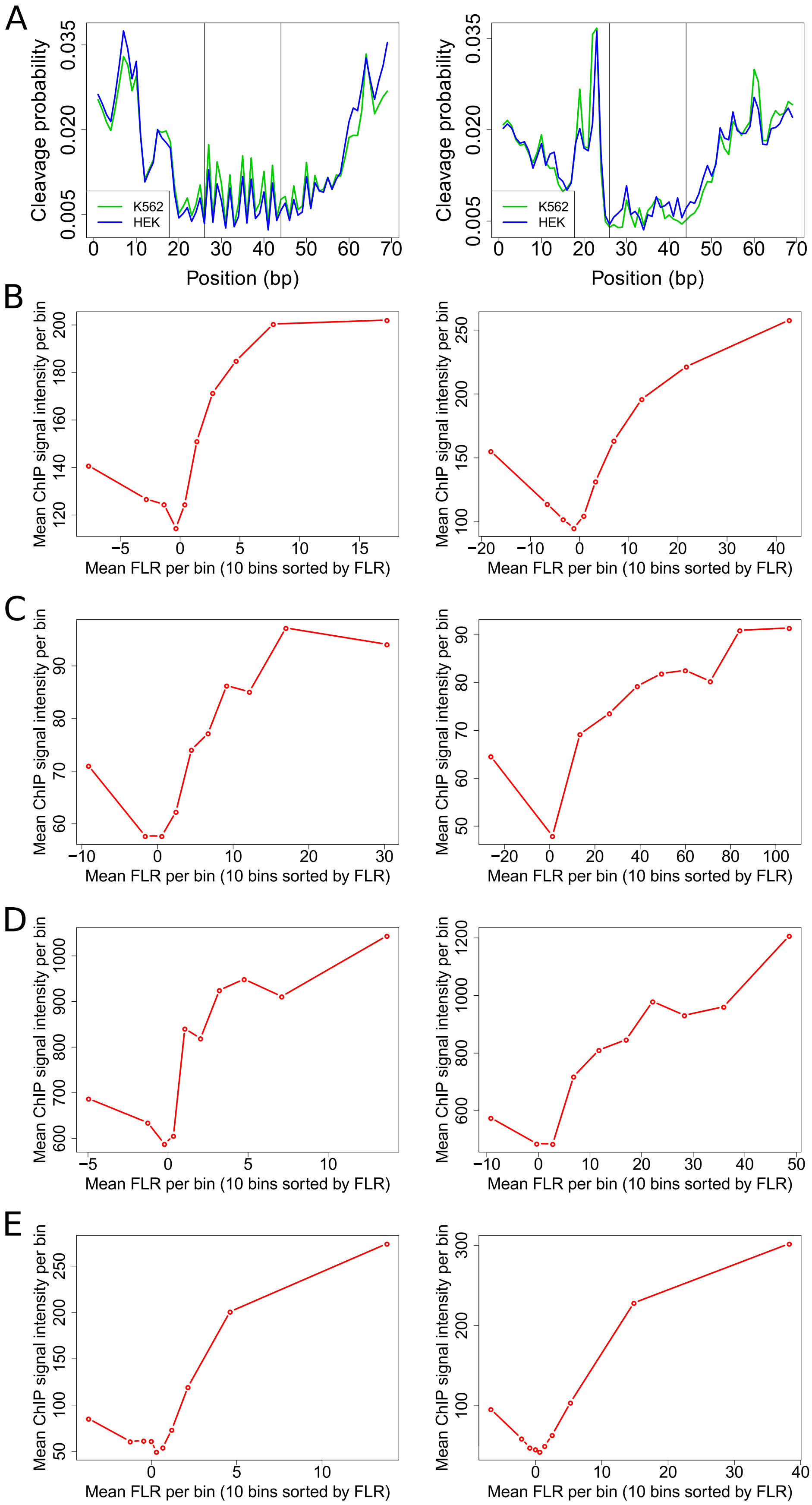


**Supplementary Figure 6:** The relevance of the learned footprint models. (A) Identical CTCF footprint profiles in HEK293 and K562 ATAC-seq (left) and DNase-seq (right) datasets. (B-E) Concordance between ChIP-seq signal intensities and footprint scores (FLR) in K562 ATAC-seq (left) and DNase-seq (right) data for (B) CTCF, (C) NRF1, (D) CREB1 and (E) USF1. Motif sites that overlap ChIP-seq peaks are divided in ten bins according to FLR. The mean ChIP-seq signal intensity and FLR is plotted for each bin.

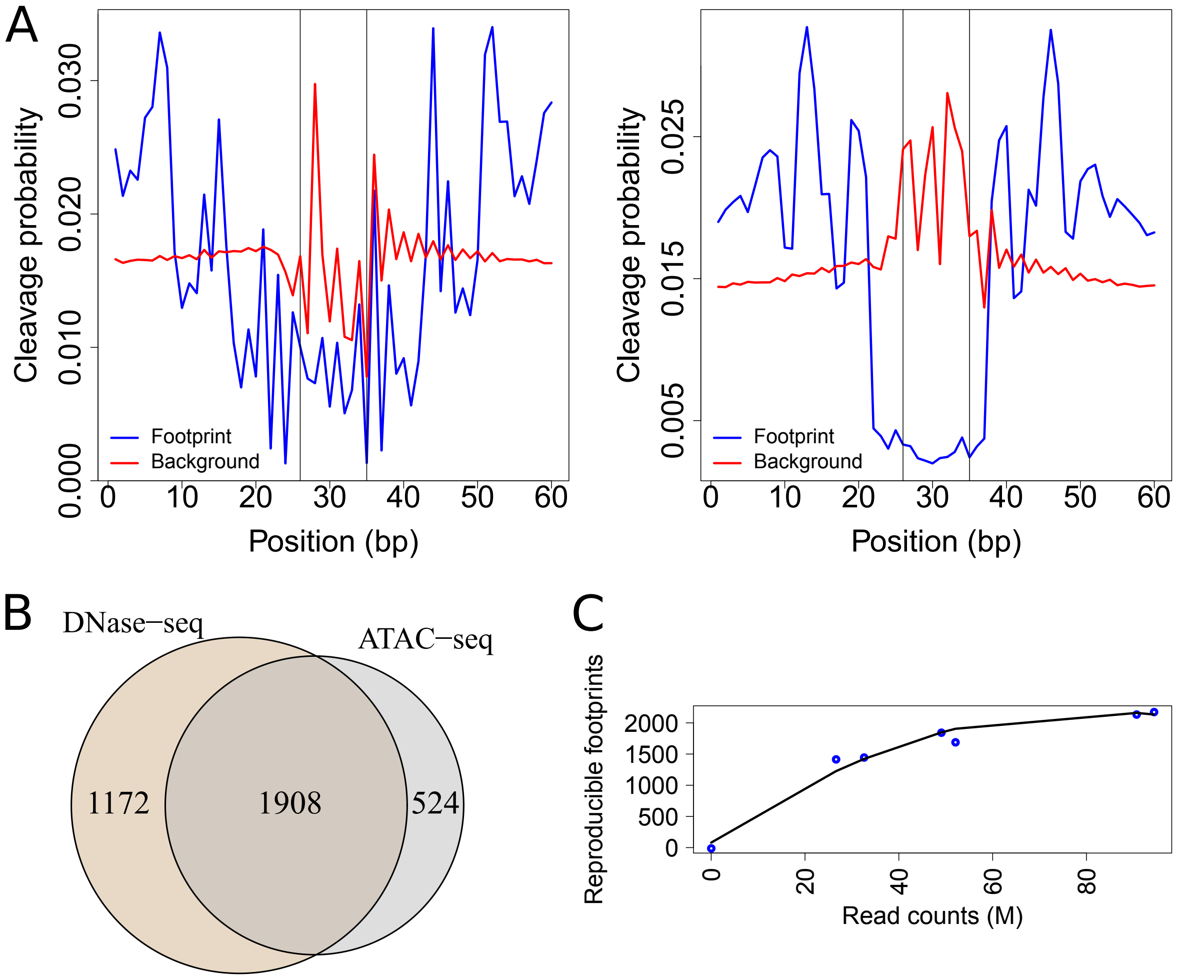


**Supplementary Figure 7:** Analysis of NRF1 footprints. (A) NRF1 footprints inferred from K562 ATAC-seq data (left) and DNase-seq data (right). Vertical lines depict the edges of the motif match. (B) Overlap between reproducible NRF1 footprints in the HEK293 DNase-seq and combined ATAC-seq replicates, found using the footprint models learned from the K562 data. (C) Numbers of reproducible NRF1 footprints in HEK293 ATAC-seq datasets at different depths.

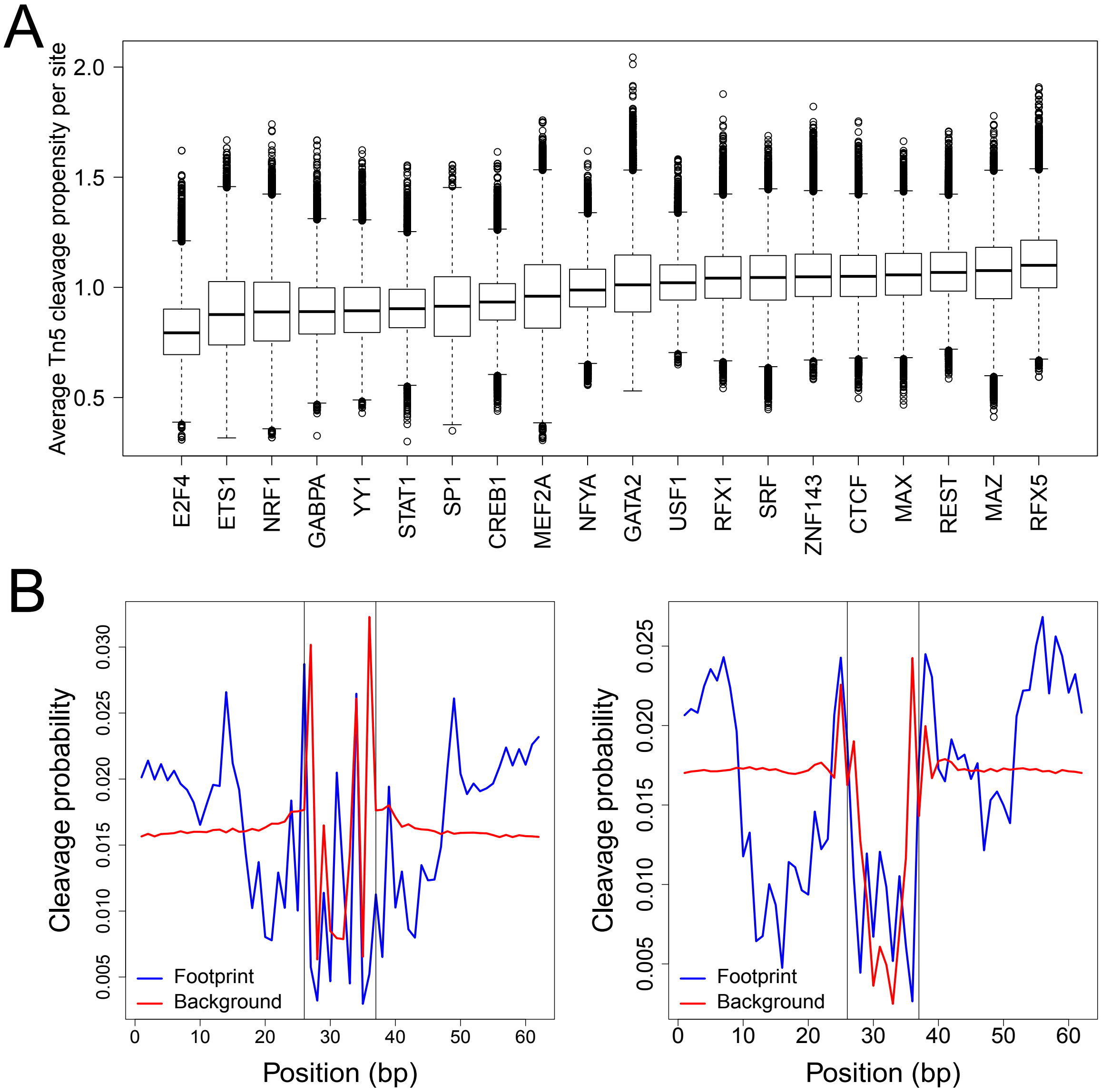


**Supplementary Figure 8:** Method and TF-specific footprinting efficiency. (A) Average Tn5 cleavage propensities over candidate TFBSs for all 20 assayed factors. (B) MEF2A footprints inferred from K562 ATAC-seq data (left) and DNase-seq data (right). Vertical lines depict the edges of the motif match.

| Cell type | Sample description | Total mapped read pairs | Percent mtDNA | Percent uniquely aligned after removing mtDNA | Percent duplication after removing mtDNA | Final read pairs after processing |
|---|---|---|---|---|---|---|
| K562 | 10 minute lysis | 98241437 | 74.9 | 60.86 | 36.1 | 11824634 |
| K562 | 5 minute lysis | 59725560 | 73.3 | 61.68 | 28.52 | 8293938 |
| K562 | No lysis buffer | 64162804 | 18 | 76.09 | 28.83 | 26203527 |
| HEK293 | High depth, bio1-tech1 | 212332636 | 21.7 | 79.15 | 38.6 | 74957855 |
| HEK293 | High depth, bio1-tech2 | 215849442 | 16.7 | 79.41 | 42.41 | 75883012 |
| HEK293 | High depth, bio2-tech1 | 189055455 | 8.3 | 80.35 | 17.42 | 106390553 |
| HEK293 | High depth, bio2-tech2 | 212178995 | 3.4 | 80.84 | 25.81 | 112909794 |
| HEK293 | Medium depth, bio1-tech1 | 101177506 | 22 | 78.93 | 22.54 | 44903594 |
| HEK293 | Medium depth, bio1-tech2 | 115293922 | 16.9 | 79.12 | 27.43 | 50914321 |
| HEK293 | Medium depth, bio2-tech1 | 85731217 | 8.4 | 80.28 | 8.82 | 53211877 |
| HEK293 | Low depth, bio1-tech1 | 53199070 | 21.9 | 78.99 | 12.83 | 26607741 |
| HEK293 | Low depth, bio1-tech2 | 59968056 | 16.8 | 79.19 | 15.84 | 30798873 |
| HEK293 | Low depth, bio2-tech1 | 40964758 | 8.4 | 80.3 | 4.54 | 26613414 |
| HEK293 | Low depth, bio2-tech2 | 51835433 | 3.4 | 80.81 | 7.72 | 34364305 |

**Supplementary table 1:** General statistics of the ATAC-seq datasets generated in the study.

| Cell type | Data type | Description | Accession code | Library depth after processing |
|---|---|---|---|---|
| K562 | DNase-seq | Replicate 1 (ENCODE) | ENCFF000SWU | 72166285 |
| K562 | DNase-seq | Replicate 2 (ENCODE) | ENCFF000SXA | 138770111 |
| K562 | DNase-seq | Replicate 3 (ENCODE) | ENCFF000SWY | 88033023 |
| K562 | DNase-seq | Replicate lab | Generated for the study | 134851555 |
| HEK293 | DNase-seq | Replicate 1 (ENCODE) | ENCFF000SPK | 68339552 |
| HEK293 | DNase-seq | Replicate 2 (ENCODE) | ENCFF000SQB | 164469299 |
| HEK293 | DNase-seq | Replicate lab | Generated for the study | 126253898 |
| Human (YH1) | Tn5 transposition | Deproteinized genomic DNA | SRX030445 | 39753928 |
| D. melanogaster | Tn5 transposition | Deproteinized genomic DNA | SRX030438 | 22705812 |

**Supplementary table 2:** Descriptions, accession codes and final read counts for the utilized DNase-seq datasets and libraries generated by Tn5 transposition of deproteinized genomic DNA.

| Comparison name | Biological replicate 1 | Biological replicate 2 |
|---|---|---|
| High depth ATAC-seq 1 | High depth, bio1-tech1 | High depth, bio2-tech1 |
| High depth ATAC-seq 2 | High depth, bio1-tech2 | High depth, bio2-tech2 |
| Medium depth ATAC-seq 1 | Medium depth, bio1-tech1 | Medium depth, bio2-tech1 |
| Medium depth ATAC-seq 2 | Medium depth, bio1-tech2 | Medium depth, bio2-tech1 |
| Low depth ATAC-seq 1 | Low depth, bio1-tech1 | Low depth, bio2-tech1 |
| Low depth ATAC-seq 1 | Low depth, bio1-tech2 | Low depth, bio2-tech2 |

**Supplementary table 3:** Scheme for ATAC-seq library comparisons for JAMM-IDR peak calls or FLR-IDR footprint calls.

| Cell line | Factor | Accession code |
|---|---|---|
| HEK293 | CTCF | ENCFF002DCV |
| K562 | CREB1 | ENCFF001UJI, ENCFF001UJJ |
| K562 | CTCF | ENCFF002CEL, ENCFF002CLS, ENCFF002CWL, ENCFF002DBD, ENCFF002DDJ |
| K562 | E2F4 | ENCFF002CWM |
| K562 | ETS1 | ENCFF002CLX |
| K562 | GABPA | ENCFF002CLZ |
| K562 | GATA2 | ENCFF002CMA, ENCFF002CWQ |
| K562 | MAX | ENCFF002CXD |
| K562 | MAZ | ENCFF002CXE |
| K562 | MEF2A | ENCFF002CMD |
| K562 | NFYA | ENCFF002CXI |
| K562 | NRF1 | ENCFF002CXK, ENCFF454OVP, ENCFF657YIC, ENCFF664FFU |
| K562 | REST | ENCFF002CMF |
| K562 | RFX1 | ENCFF654RTP |
| K562 | RFX5 | ENCFF002CXV |
| K562 | SP1 | ENCFF002CMN, ENCFF191QSX |
| K562 | SRF | ENCFF002CMP |
| K562 | STAT1 | ENCFF002CYB, ENCFF002CYC, ENCFF002CYD, ENCFF002CYE |
| K562 | USF1 | ENCFF002CMV |
| K562 | YY1 | ENCFF002CMW, ENCFF002CMX, ENCFF002CYQ |
| K562 | ZNF143 | ENCFF002CYR |

**Supplementary table 4:** ChIP-seq peaks used in the analysis.

| Factor | PWM |
|---|---|
| CREB1 | MA0018.2 |
| CTCF | MA0139.1 |
| E2F4 | M5180_1.01 |
| ETS1 | MA0098.1 |
| GABPA | MA0062.2 |
| GATA2 | MA0036.1 |
| MAX | M5613_1.02 |
| MAZ | M00649 |
| MEF2A | M5615_1.02 |
| NFYA | MA0060.1 |
| NRF1 | M00652 |
| REST | MA0138.2 |
| RFX1 | M00280 |
| RFX5 | M5779_1.02 |
| SP1 | MA0079.2 |
| SRF | MA0083.1 |
| STAT1 | MA0137.2 |
| USF1 | M5943_1.02 |
| YY1 | M5954_1.02 |
| ZNF143 | M5966_1.02 |

**Supplementary table 5:** PWM IDs used for genome-wide motif searches.
